# A method for achieving complete microbial genomes and improving bins from metagenomics data

**DOI:** 10.1101/2020.03.05.979740

**Authors:** Lauren M. Lui, Torben N. Nielsen, Adam P. Arkin

**Affiliations:** Environmental Genomics and Systems Biology Division, Lawrence Berkeley National Laboratory, Berkeley, CA, USA; Department of Bioengineering, University of California, Berkeley, CA, USA; Innovative Genomics Institute, Berkeley, CA, USA

## Abstract

Metagenomics facilitates the study of the genetic information from uncultured microbes and complex microbial communities. Assembling complete microbial genomes (*i.e*., circular with no misassemblies) from metagenomics data is difficult because most samples have high organismal complexity and strain diversity. Less than 100 circularized bacterial and archaeal genomes have been assembled from metagenomics data despite the thousands of datasets that are available. Circularized genomes are important for (1) building a reference collection as scaffolds for future assemblies, (2) providing complete gene content of a genome, (3) confirming little or no contamination of a genome, (4) studying the genomic context and synteny of genes, and (5) linking protein coding genes to ribosomal RNA genes to aid metabolic inference in 16S rRNA gene sequencing studies. We developed a method to achieve circularized genomes using iterative assembly, binning, and read mapping. In addition, this method exposes potential misassemblies from k-mer based assemblies. We chose species of the Candidate Phyla Radiation (CPR) to focus our initial efforts because they have small genomes and are only known to have one ribosomal RNA operon. We present 34 circular CPR genomes, one circular Margulisbacteria genome, and two circular megaphage genomes from 19 public and published datasets. We demonstrate findings that would likely be difficult without circularizing genomes, including that ribosomal genes are likely not operonic in the majority of CPR, and that some CPR harbor diverged forms of RNase P RNA. Code and a tutorial for this method is available at https://github.com/lmlui/Jorg.

## Introduction

Shotgun metagenomics and marker gene sequencing are powerful tools to survey and study organisms that we cannot yet isolate and culture in the laboratory. This is especially true for environmental samples where culturability estimates for bacterial and archaeal communities range from ∼22-53% for soil, ∼10-70% for ocean and lakes, and ∼8-32% for ocean sediment [1]. Scientists have turned to shotgun metagenomics to provide genome-resolved analysis of complex samples, but assembling genomes from shotgun metagenomics data is inherently more difficult than assembling those from cultured isolates. Challenges in metagenomics assembly arise from the heterogeneity of samples, available sequencing technology, and the limitations of bioinformatics algorithms we use for assembly and genome binning. Metagenomes contain uneven amounts of an unknown number of genomes, which creates a compounded computational problem in terms of simplifying assumptions, time, and computer memory.

In the 1990s when the first genomes were sequenced and assembled, scientists used long reads from Sanger sequencing and overlap layout consensus (OLC) methods for assembly [2]. With the development of next-generation sequencing technologies, we gained the ability to sequence millions of reads at a massively reduced cost, but using traditional OLC algorithms became too computationally intensive. The computational complexity of OLC algorithms scale as the square of the number of input reads (because each read is compared to every other read), so they are impractical for datasets of millions of reads, compared to the thousands of reads generated from Sanger sequencing. To handle the deluge of sequencing data (in terms of the volume of reads and projects) de Bruijn graph assembly methods were developed. The time and memory complexity of de Bruijn based assembly algorithms typically scale with the size of the metagenome instead of the number of reads. However, de Bruijn graph methods can introduce misassemblies; due to the decomposition of reads into k-mers, context is lost and it is possible for the graphs to contain paths that do not correspond to real genomic sequence [3,4]. Traditional OLC assemblers such as the Celera Assembler [5], SGA [6] and MIRA [7] ensure that only contigs consistent with actual genome sequence are produced (this is sometimes referred to as maintaining read coherence). Using long-read sequencing can overcome some of the issues with k-mer based assembly and newer assemblers for this type of data have started using OLC assembly methods again [3]. However, long-read sequencing requires much larger amounts of DNA (micrograms) that is high quality and high molecular weight (10-50kb) as compared to short read technologies (as little as 1 ng). Extracting enough high molecular weight DNA can be limited by sampling costs and available biomass, especially for certain types of environmental samples, so short read sequencing may sometimes be the only option.

Beyond assembly, a key challenge with metagenomics is grouping contigs into genome bins. We use “contig” in the way it was originally defined by Rodger Staden, where a contig is a set of overlapping segments of DNA from shotgun sequencing [8]. It is rare for a complete genome to be assembled into a single piece *de novo* from short reads, so contigs are grouped into “bins,” often based on coverage and tetranucleotide frequencies. If two contigs belong to the same genome, they are expected to have similar coverage and tetranucleotide profiles [9]. However, coverage has problems for multiple reasons. If a particular microbe is growing rapidly, some regions may have higher coverage than the rest of the genome [10]. In addition, for organisms where the copy number of ribosomal RNA (rRNA) operons exceeds unity, the contig(s) with the rRNA genes will not have the same coverage as the rest of the contigs in the genome. This is also true of other multi-copy genes and other repetitive elements. Tetranucleotide frequencies are problematic because horizontally transferred regions may have different frequencies than the rest of the genome [11] and this can result in such pieces being put into different bins by the binning algorithm. Despite these issues, binning is helpful in identifying potential genomes in metagenomics data, especially when using short read sequencing technologies.

To evaluate the quality of a bin, the metrics of “contamination” and “completeness” are often used. Completeness and contamination are detected generally by looking for violations of conserved features of complete isolate genomes. Such features include having complete sets of universally (or at least phylogenetically) conserved single-copy protein genes without any duplication or excessive variation in tetranucleotide frequency. Other measures of completeness have been suggested, such as establishment of a core conserved set of ubiquitous genes. Tools such as RefineM [12] and CheckM [13] apply these rules to assemblies to determine completeness and contamination. However, these tools are not always accurate for species that are not well studied. Candidate Phyla Radiation (CPR) species are often classified as having 60-80% completeness by these tools, even for circular genomes. To overcome challenges of binning, scientists have started to assemble circular, complete genomes from metagenomes [14–23], which are also called CMAGs (complete metagenomic-assembled genomes) [24]. In comparison to genome bins, a high quality reference collection that controls for misassemblies and is composed of circular genomes (1) provides more accurate inference of identity and estimation of capabilities of uncultured microbes within complex microbiomes, (2) allows more accurate taxonomic assessment of the composition of these microbiomes through better linkage of marker genes in single organisms, (3) provides high-quality scaffolds on which reads can be assembled, both to allow measures of strain variation within a microbiome study and to aid in better assembly of reads across many microbiome samples, and (4) affords the ability to study synteny and genomic context of genes in these organisms. In addition, while there are existing methods for generating high-quality MAGs, there is evidence that these MAGs still contain significant contamination by exogenous sequence and have misassemblies triggered by lack of read coherence. Circularization of genomes helps assure that there is likely no contamination in the assemblies. Despite the advantages of circularizing genomes, very few metagenomics studies to date (<30) have published circular genomes [14,15,24,16–23].

We describe a semi-automated method that facilitates recovery of circular archaeal and bacterial genomes from metagenomics data and that also provides checks for misassemblies. In this study we do not focus on how many circularized genomes we can get from a dataset, but rather the method itself to help circularize bins of interest and to ensure that they are high quality. To assist with the travails of circularizing genomes, our method overcomes issues from using k-mer based assembly and automates iterative extension of contigs. Our general approach is to produce a “standard” metagenomic assembly, bin using a “standard” binning tool, extract reads based on k-mer similarity and reassemble these using a “standard” isolate-focused assembler. To demonstrate this method, we have obtained 34 circular CPR genomes, one circular Margulisbacteria genome, and two circular megaphage genomes from 19 public and published metagenomics datasets. To our knowledge, only 41 other CPR circularized genomes have been published from 9 studies [14–22], so we believe this to be the largest presentation of circularized CPR genomes in a single study. With this set, we demonstrate findings that would likely be difficult without a large number of unique circularized genomes, including that ribosomal genes are likely not operonic in the majority of CPR and finding diverged forms of RNase P RNA in CPR species.

## Results

### Circularization Method

To have confidence that the genomes we generate match real organisms, we looked for criteria that would indicate that a genome is circular and complete. The literature is replete with techniques and proposals for measuring the completeness of genomes and to what level they are complete [13,16], but these often have difficulty when encountering novel genomes because their criteria are based on known isolate genomes. We focus on evidence for incompleteness in terms of missing essential genes that are found across the tree of life. That is, we are more concerned with ensuring that anything we label a circularized genome meets basic criteria that indicates that it is not incomplete. We posit that a complete, circularized genome must satisfy at least the following conditions:

1. The genome is either circular or there is solid evidence that it is linear. While rare, linear bacterial genomes exist [25].
2. The genome has a full complement of rRNAs (16S, 23S, 5S), transfer RNAs (all amino acids represented), and RNase P RNA (since this is nearly universally necessary to process tRNA transcripts). Absence of any of these genes must be explained. We advocate using these as a check for a complete genome instead of single copy marker protein genes because checks for single copy marker protein genes can vary by clade; in only rare instances would these noncoding RNA genes be missing [26,27].
3. There is significant read coverage across the entire genome. Assemblies that rely on single reads for continuity are prone to error. With some exceptions for very high coverage organisms, we generally require minimum coverage no lower than 30% of the average coverage.

To develop and test this method, we mined the Sequence Read Archive (SRA) hosted at the National Center for Biotechnology Information (NCBI) [28] for metagenomic sequencing data generated from groundwater samples, where CPR are prevalent (see Table 1 for a list of the datasets used in this study. We focused on assembling CPR genomes because they are (1) small and thus easier to assemble, (2) to the best of our knowledge only have one set of rRNA genes. These two criteria gave us the easiest targets for circularization.

**Table 1:** Description of Metagenomes in this Study. We focused on groundwater datasets because they have a higher fraction of CPR. Many of the datasets are from studies of anthropologically contaminated sites. All identifiers are SRA except for SURF_D.

| Identifiers | Study Description | Reference |
| --- | --- | --- |
| ERX2165959 | Groundwater from monitoring wells from naphthalene contaminated surface sediments, where effluent from the coal-tar contaminated groundwater surfaces | [29] |
| SRX1085364 | Terrestrial subsurface C, N, S and H cycles cross-linked by metabolic handoffs | [16] |
| SRX1775573<br>SRX1775577<br>SRX1775579 | Potential for microbial H<sub>2</sub> and metal transformations associated with novel bacteria and archaea in deep terrestrial subsurface sediments | [21] |
| SRX1990955 | Groundwater microbial communities from Rifle, Colorado - Rifle Oxygen_injection A2 metagenome | [16] |
| SRX2838984 | Coupling Microbial Communities to Carbon and Contaminant Biogeochemistry in the Groundwater-Surface Water Interaction Zone | [30–32] |
| SRX3024504<br>SRX3024507<br>SRX3024508 | DNA from groundwater after nitrate injection, filter size 0.2µm and 0.1µm | [33] |
| SRX3307784 | Subsurface groundwater microbial communities from S. Glens Falls, New York, USA - GMW37 contaminated, 5.8 m metagenome | [29] |
| SRX3348993 | Development of a pipeline for high-throughput recovery of near-complete and complete microbial genomes from complex metagenomic datasets: Groundwater sample from aquifer - Crystal Geyser CG19_WC_8/21/14_NA | [34] |
| SRX3574179 | Investigating microbial roles in methane emission, contaminant degradation, and biogeochemical cycles in an aquifer near a municipal landfill | Laura Hug Lab;<br>https://uwaterloo.ca/hug-research-group/ |
| SRX3602289<br>SRX3602720 | Groundwater microbial communities from the Aspo Hard Rock Laboratory (HRL) deep subsurface site, Sweden | Mark Dopson Lab;<br>https://lnu.se/en/staff/mark.dopson/ |
| SURF_D | Groundwater samples from the Sanford Underground Research Facility (SURF) | [35] |

The first steps of the method are standard to regular metagenomics assembly pipelines. For each metagenome, we trimmed the reads to remove any remaining Illumina adapter fragments and low quality ends, as well as whole reads that weren’t of sufficient quality, using BBtools (Fig 1A). Next, we assembled the processed reads using SPAdes [36] (Fig 1B). We proceeded with successful assemblies and used MetaBAT 2 [37] (Fig 1C) to produce a collection of bins for each. We went through 188 assembled metagenomes and picked bins with 5 or fewer contigs and coverage above 40X, although we made exceptions for bins that looked promising, such as a bin with many contigs, but with one or two large contigs that comprise most of the bin’s sequence length (Table 2). We used GTDB-Tk [38] to classify the bins and picked a set of CPR bins. We used these bins as “bait” to select read pairs for use with the isolate-focused assembler MIRA (Fig 1D).

**Table 2:** List of 35 circularized bacterial genomes in this study. Thirty-four of the genomes are classified as CPR and one is classified as a Margulisbacteria by GTDB-Tk. The coverage, original number of contigs, and length of the original bin is also included.

| Genome ID | GTDB-TK<br>Taxonomy<br>(Class) | Genome<br>Size (bp) | Original Bin Stats |  |  |
| --- | --- | --- | --- | --- | --- |
|  |  |  | Num of<br>Contigs | Coverage | Size of<br>Bin (bp) |
| ERX2165959_bin_184 | Paceibacteria | 523910 | 2 | 126X | 531828 |
| ERX2165959_bin_23 | Microgenomatia | 1147419 | 1 | 134X | 1146985 |
| ERX2165959_bin_53 | Microgenomatia | 646630 | 3 | 116X | 780569 |
| ERX2165959_bin_80 | Microgenomatia | 1014979 | 1 | 216X | 1024366 |
| SRX1085364_bin_95 | Microgenomatia | 819458 | 4 | 109X | 826172 |
| SRX1775573_bin_5 | Gracilibacteria | 998919 | 1 | 104X | 999239 |
| SRX1775577_bin_36 | Gracilibacteria | 999108 | 1 | 74X | 998727 |
| SRX1775579_bin_0 | Dojkabacteria | 732899 | 1 | 51X | 733907 |
| SRX1990955_bin_0 | Margulisbacteria<br>(phylum); WOR-1 | 1676518 | 1 | 56X | 1673447 |
| SRX1990959_bin_38 | Paceibacteria | 585024 | 2 | 35X | 582583 |
| SRX2838984_bin_5 | Paceibacteria | 1030062 | 7 | 274X | 1030337 |
| SRX3024504_bin_47 | Paceibacteria | 672946 | 1 | 41X | 673298 |
| SRX3024507_bin_14 | ABY1 | 1064268 | 4 | 48X | 1106392 |
| SRX3024507_bin_96 | Paceibacteria | 672946 | 1 | 29X | 673073 |
| SRX3024508_bin_27 | Paceibacteria | 672946 | 1 | 91X | 673184 |
| SRX3307784_bin_186 | Paceibacteria | 581622 | 4 | 174X | 578873 |
| SRX3307784_bin_197 | Paceibacteria | 822324 | 8 | 203X | 962091 |
| SRX3307784_bin_224 | Microgenomatia | 646579 | 3 | 118X | 780569 |
| SRX3307784_bin_45 | UBA1384 | 872881 | 1 | 93X | 872947 |
| SRX3307784_bin_80 | Microgenomatia | 1013439 | 1 | 220X | 1024366 |
| SRX3307784_bin_91 | Paceibacteria; | 523446 | 3 | 129X | 531688 |
| SRX3348993_bin_93 | Saccharimonadia | 1005778 | 1 | 64X | 1005352 |
| SRX3574179_bin_116 | Saccharimonadia | 949592 | 4 | 37X | 1000785 |
| SRX3574179_bin_12 | ABY1 | 1027227 | 1 | 135X | 1028393 |
| SRX3574179_bin_242 | Microgenomatia | 1127729 | 3 | 48X | 1130277 |
| SRX3574179_bin_244 | Paceibacteria | 707009 | 1 | 106X | 708095 |
| SRX3574179_bin_38 | Paceibacteria | 682291 | 5 | 124X | 680516 |
| SRX3574179_bin_63 | ABY1 | 1209998 | 2 | 88X | 1209921 |
| SRX3574179_bin_75 | ABY1 | 954552 | 3 | 152X | 979051 |
| SRX3602289_bin_51 | Microgenomatia | 852061 | 2 | 94X | 853126 |
| SRX3602720_bin_127 | Paceibacteria | 750123 | 18 | 160X | 826132 |
| SRX3602720_bin_74 | ABY1 | 1035066 | 3 | 32X | 1041342 |
| SRX5650846_bin_20 | Microgenomatia | 947771 | 9 | 53X | 949620 |
| SURF_D_bin_21 | Microgenomatia | 978848 | 3 | 193X | 978553 |
| SURF_D_bin_31 | Microgenomatia | 1496927 | 5 | 159X | 1501138 |

**Fig 1.**
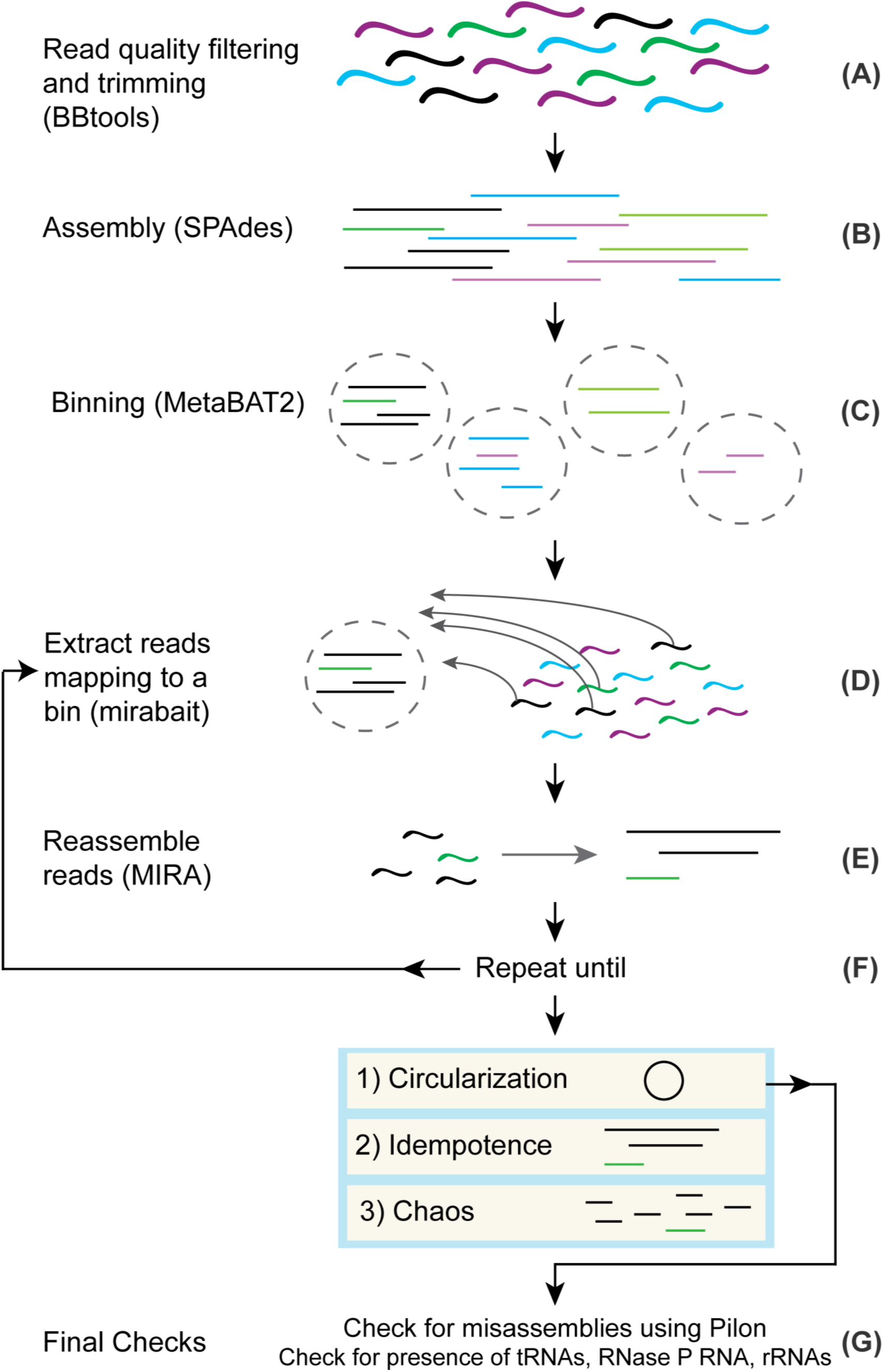
General method for circularizing genomes from metagenomes. (A) Reads have adapters trimmed and low quality reads are filtered using BBtools. (B) Processed reads are assembled into contigs using SPAdes. (C) Contigs are grouped into bins using MetaBAT 2. (D) After choosing a bin for circularization, reads mapping to the bin are extracted from the original processed reads and used as input into (E) where they are assembled into contigs using MIRA. (F) Steps D and E are repeated as necessary until the bin is deemed to be in a Circularization, Idempotence, or Chaos state. (G) If a bin is deemed circular, we do final checks for misassemblies using Pilon and for the presence of rRNA, tRNAs, and RNase P RNA before officially calling the bin a circularized genome.

The purpose for assembling contigs first with SPAdes and then switching to MIRA with a subset of reads is that the computational requirements of MIRA make it impractical as a metagenome assembler. This is in part because MIRA does full alignment of the reads during assembly. We would like to note that OLC metagenomics assemblers exist [3], but their memory and time requirements are high compared to SPAdes or are not appropriate for assembling bacterial genomes with paired-end sequencing data. MIRA has been used to extract mitochondrial genomes [39] from eukaryotic sequencing projects. Our approach is very similar; instead of providing seed sequences to separate the mitochondrial genomes from the eukaryotic DNA, we use bins as seeds to separate genomes from the entire metagenomics dataset. In our experience, MIRA produces superior results for isolates, and it also provides additional features that benefit our method. MIRA comes with the tool mirabait, which provides support for extracting read pairs based on k-mer content. MIRA also has a variety of features that help expose problematic parts of assemblies. For example, MIRA sets tags to indicate parts of the assembly that may require manual intervention, based on changes in coverage, GC, and other anomalies. These tags are extremely useful in conjunction with traditional assembly finishing tools such as Gap4/Gap5 [40].

Perhaps even more critical to this method than MIRA’s utilities is the fact that MIRA also ensures read coherence as an overlap-based assembler, unlike k-mer based assemblers like SPAdes. SPAdes is commonly used for metagenomics assembly, and in our experience, produces results that are as good as any other metagenomics assembler that is typically recommended [4]. However, there are often misassemblies caused by running SPAdes on a large and heterogeneous collection of metagenomes with the same set of k-mers. Ideally, the user would conduct tests to find the optimal collection of k-mers for each individual metagenome, but this step is time consuming. Thus, many users - us included - pick a canonical set like 21, 33, 55, 77, 99 and 127 that in most cases give the greatest contiguity in the assembly. Unfortunately, this practice can produce illusionary contiguity if the read coverage cannot support all of the k-mer sizes [3]. Larger k-mers increase contiguity, but the read coverage may not support them. By using MIRA, contigs that do not have read coherence may be exposed.

After we used mirabait to extract read pairs that mapped to selected bins (Fig 1D) and reassembled them using MIRA (Fig 1E), we iterated these two steps (Fig 1F). This iterative process results in “digital primer walking” to extend the contigs of the bin, similar to primer or genome walking that was initially used to sequence genomes in the late 1980s to early 1990s [41]. At each iteration, reads with a portion mapping to any part of a contig will be included and can lead to extension or fusion of contigs. We specifically chose to reassemble all of the reads during each iteration to provide a more robust handling of repeats. On occasion, the extension of the contigs resulted in overlap with contigs from other bins and unbinned contigs. Manually including these contigs as part of the bait can speed up the process significantly. However, we also routinely examined intermediate results and, in some cases, we saw anomalous coverage values for different contigs indicating possible chimerism. If we saw the bin containing contigs with significantly different coverage values (>10% difference), we removed the offending contigs and restarted. We iterated this process until one of these outcomes occurred:

1. **Circularization**. For us to decide that this had occurred, we looked for a single contig with a significant - and exact - repeat at the ends. In addition, we required that the repeat be at least 100 nt in length, was longer than any other repeat in the contig, and did not match any of the other repeats.
2. **Idempotence**. In some cases, we observed no change in the assembled contigs after a round of read pair extraction and reassembly with MIRA. We examined some of these instances in detail and we believe the change in coverage causes MIRA to refuse to continue extending contigs. It is possible to adjust MIRA’s thresholds of what constitutes low and high coverage to allow contig extension to continue. However, this modification increases the risk of collapsing repeats or creating chimeric assemblies.
3. **Chaos**. There are cases where a bin is shattered into a multitude of pieces. We are not certain as to the exact cause, but this result is likely due to misassemblies from the initial SPAdes assembly (discussed in more depth in a later section). Chaos appears strongly correlated with GC and tends to occur more often when the GC content is high. We have investigated a few in more detail and for some found that the contigs that shatter have low 127-mer coverage as reported by SPAdes. We believe that Chaos bins are caused by lack of read coherence in the contigs and if that is indeed the case, there is little we can do. Once we see Chaos set in, it appears to be permanent.

After circularizing a contig, we did final checks for misassemblies with Pilon (Fig 1G). We used Pilon [42] on the contig and then we rotated it by half the length to ensure that the ends were in the middle and applied Pilon again (Fig 1G). We rotate the genomes because Pilon is not capable of covering the ends of a contig. While Pilon found minor insertions/deletions due to the circularization, it did not find any other issues in the genomes.

We next searched the genomes for a full complement of ribosomal RNAs (16S, 23S, 5S), tRNAs (all amino acids represented) and RNase P RNA to check that the genome was correctly circularized and was not missing regions. For RNase P RNA, we needed to manually reduce score thresholds to find all RNase P RNAs (discussed in more detail in a later section). We were able to find tRNAs for all amino acids, although some tRNAs had Group I introns, making them difficult to detect. Structural RNAs are sometimes invaded by Group I introns, which is particularly true for tRNAs [Patricia Chan, private communication]. When a genome passed the final check with the detection of the set of non-coding RNAs, we considered the genome to be accurately circularized.

SRX3307784_bin_197 is an example of a bin that appeared to be circular, but did not pass the check of having RNase P RNA. The assembled contig had a solid 414 base pairs of overlap at the ends, ribosomal proteins present and tRNAs for all amino acids. However, we did

not find a copy of RNase P RNA even when we lowered the detection threshold. This caused us to look closer and we discovered that there was another contig in the assembly which we had thought was contamination after the initial circularization. This contig has a copy of RNase P RNA and we were able to incorporate it into the assembly after we discovered a repeat that was too long for the reads to span and that Pilon did not detect. We came to the conclusion that this was a case of false circularization. To address the misassembly, we put the bin through more iterations with mirabait and MIRA, which resulted in a larger genome which passed all of the final checks.

In approximately 10% of the cases we attempted, we succeeded in creating a circularized genome out of a bin. However, our selection of bins was not random as we were heavily biased in favor of bins that we judged easiest to circularize, such as selecting bins classified as CPR, had relatively few contigs, and had solid coverage. We are confident that we can assemble more genomes from these datasets because we have also been able to circularize genomes from archaea and other bacteria; these genomes will be published in future papers. We intend to make these genomes generally accessible as we finish them. The code for iterating to pull reads mapping to a bin and reassembly with MIRA are available on Github (https://github.com/lmlui/Jorg) and we are currently making an app in the Department of Energy KnowledgeBase [43].

### Description of circularized CPR genomes

Using our method, we circularized 34 CPR genomes and one Margulisbacteria genome (Table 2 and Fig 2). To create a phylogenetic tree, we used a structural alignment 16S rRNA genes. During this process we found that many of the 16S included large introns with LAGLIDADG homing endonucleases, an observation that has been noted in other CPR studies [17].

**Fig 2.**
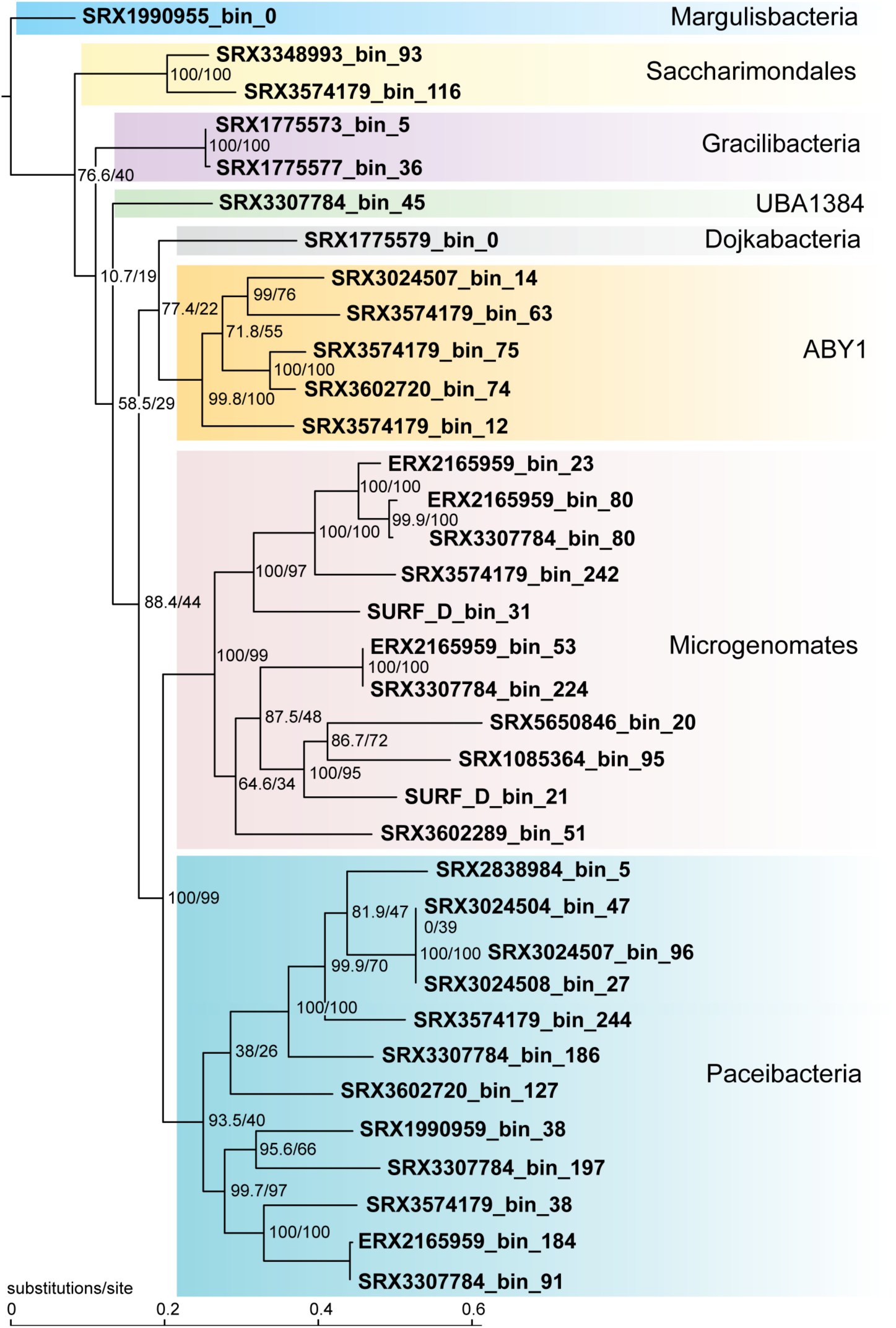
Phylogenetic Tree based on SSU structural alignment. At the base of the branches, in the fraction the top value is SH-aLRT which is a branch test [44] and the bottom value is the bootstrap value. Class is listed for the CPR based on GTDB-TK taxonomy of that genome. The class UBA1384 is also known as Berkelbacteria. We specified the Margulisbacteria as the outgroup when creating the tree using IQ-TREE [45].

In general, these genomes are novel, but in one case, SRX1085364_bin_95, we found that the genome is 100% identical to a previously circularized genome (INSDC Accession CP011214.1) from that dataset [17]. Assembling the same genome as another group helps validate our findings and that with careful manual curation two different groups can come to the same assembly result despite differences with assemblers. Four of the other genomes had 16S genes that had 100% hits in NCBI. Some of the 16S genes only had percent similarity in the low 80s to other sequences in Genbank.

We compared the genome sizes before and after circularization, and in most cases the size of the genome decreased after circularization compared to the original bin. Typically the genome shrank from a few hundred bases to a few thousand, but in some cases the genome shrank by more than 130kbp (Table 2). This shrinkage may be a result of SPAdes artifacts that MIRA determines to be lacking in read coherence. We are also aware of cases where SPAdes generates contigs that are effectively duplicates of each other apart from short stretches at the ends, and MIRA is able to resolve these into one contig.

### Misassembled contigs can be found with MIRA, *i.e*. Chaos

In some cases, when we attempt the reassembly step of a bin with MIRA, we end up with many more short contigs than what was in the original bin. SRX3024505_bin_48 started with just 7 contigs with coverage 21X and a GC content of 59%. Superficially, it looks like a reasonable bin. GTDB-Tk classifies it as a CPR in the Gracilibacteria class. However, after going through 5 rounds of our method, we ended up with 136 contigs, *i.e*., this a Chaos bin.

We do not know exactly what happened in the case of SRX3024505_bin_48, but we see Chaos routinely during reassembly with MIRA. In some cases, we have been able to conclude that Chaos results from insufficient read support for the largest k-mer used in the original SPAdes assembly. Put differently, the assembly graph wasn’t sufficiently well connected at the highest k-mer used. To determine if the Chaos of SRX3024505_bin_48 was solely a result of using MIRA, we used the same reads that we gave to MIRA as input into a SPAdes assembly. We ended up with 47 contigs, which was still significantly worse than the original bin. It is worth noting that the size of SRX3024505_bin_48 remained relatively constant during the testing and reassembly process. Although Chaos is a disappointing result in assembly, knowing that a bin likely has misassembled contigs is valuable.

Chaos predominantly occurs when the coverage is less than ∼30X. Most of the genomes we successfully circularized have much higher coverage. Based on our experience, we believe that coverage requirements for successful circularization of genomes from metagenomes are significantly higher than for isolates.

### Likely nearly all CPR have unlinked ribosomal operons

Typically in bacteria and archaea, the 16S, 23S, and 5S ribosomal RNA genes are found in an operon in the order 16S-23S-5S [46] (Fig 3A). In contrast, we noted that in the CPR genomes that we circularized, nearly 80% of them had unlinked 16S and 23S genes and sometimes unlinked 23S and 5S genes (27/34 genomes). We observed the following types of ribosomal operons in our circularized genomes: (1) operonic, but the 16S and 23S are separated by tRNAs on the same strand (Fig 3B), (2) operonic, but the 16S and 23S (or 23S and 5S) are separated by tRNAs and/or protein coding genes on the same strand (Fig 3C), (3) unlinked by distance, all three ribosomal rRNA genes are on the same strand but the 16S is separated from the 23S-5S or all three are separated by more than 2000bp and there are no possible intervening genes in the spacer regions that could connect the ribosomal genes in an operon (Fig 3D), and (4) unlinked because the 16S is on the opposite strand from the 23S and 5S (Fig 3E). In three cases, tRNA genes and/or protein coding genes on the same strand were located between the 16S and 23S or between the 23S and 5S, but there are 300-500bp regions between the genes, so in these cases the ribosomal genes may be uncoupled, but conservatively we counted them as operonic. In-depth analysis of gene spacing in operons of these genomes would be required to determine if these cases are operonic or not. In the SRX1085364_bin_95 genome, we noted that there is a homing endonuclease between both the 16S and 23S, and the 23S and 5S. Given that our genomes span a large part of the CPR phylogeny, we infer that most of the CPR likely have unlinked ribosomal operons.

**Fig 3.**
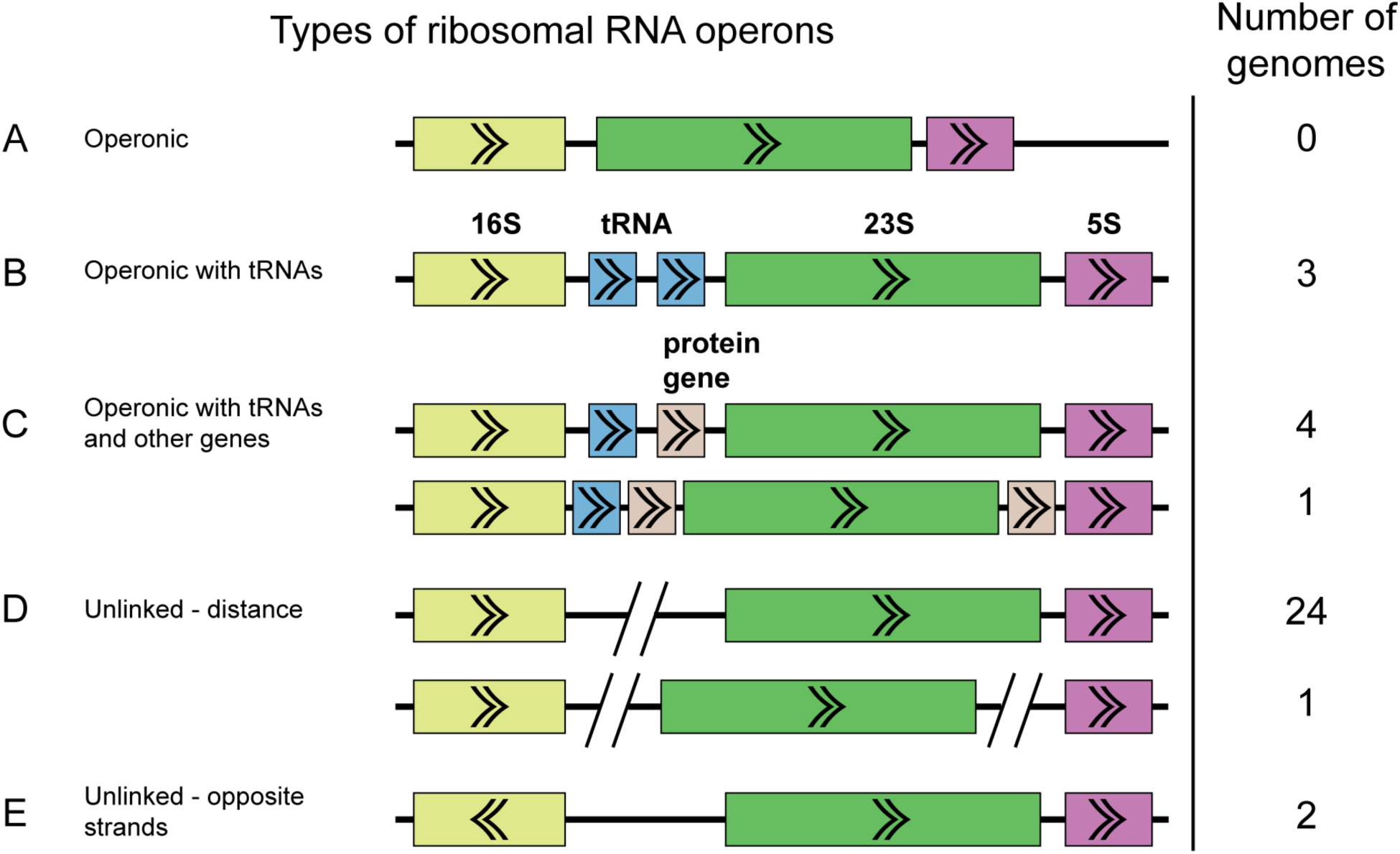
Diagram of placement types of ribosomal RNA genes. Number of genomes in this study for each category are indicated in the rightmost column. (A) Operonic. The 16S (yellow), 23S (green), and 5S (purple) ribosomal RNA genes are in an operon. (B) Operonic with tRNAs (blue). The three ribosomal RNA genes are still in an operon, but one or more tRNAs are located in the spacer between the 16S and 23S genes. (C) Operonic with tRNAs and protein coding genes (beige). The three ribosomal RNA genes are still in an operon, but one or more tRNAs or protein coding genes are located in the spacer between the 16S and 23S genes or 23S and 5S genes. It is not unusual to find that the protein coding gene is a homing endonuclease. (D) Unlinked ribosomal RNA genes by distance. The 16S gene is unlinked from the 23S and 5S genes, or the 23S is also unlinked from the 5S gene, by enough distance (>2000bp) and intervening genes on the opposite strand where it is not possible for them to be transcribed from the same promoter. (E) Unlinked ribosomal RNA genes that are on opposite strands. The 16S is on the opposite strand from the 23S and 5S genes.

The most common type of bacterial rRNA operons are those where 16S-23S-5S are transcribed together (Fig 3A), sometimes with tRNAs between the 16S and 23S (Fig 3B), so it is notable that most of the genomes in this study have unlinked rRNA operons. Instances where 16S and 23S are decoupled are unusual, although not unknown [46,47]. Separation of 23S and 5S is very unusual in bacteria but typical in archaea [47]. Decoupling between the 16S and 23S is known to occur especially in bacteria and archaea with reduced genomes (<2Mb) [46] such as *Mycoplasma gallisepticum* [48], *Borrelia burgdorferi* [49], *Ferroplasma acidarmanus* [50], as well as obligate symbionts with small genomes such as *Buchnera aphidicola* [51], *Wolbachia pipientis* [52], and *Nanoarchaeaum equitans* [47]. In a recent study of isolate genomes and pairing long reads with metagenomics data, others have also noted that a large percentage of the CPR likely have unlinked ribosomal operons based on analyzing the distance between the 16S and 23S genes [53]. However, to our knowledge, no one else has checked for tRNAs and protein coding genes comprising the operon in this type of analysis. We also do not know of other studies of CPR that have documented possible separation of 23S and 5S genes, proteins in the spacer regions between ribosomal RNA genes, and 16S and 23S on opposite strands.

### Diverged forms of RNase P RNA in CPR

RNase P is an RNA-protein endonuclease involved in the maturation of tRNAs by trimming the 5’ leader of pre-tRNAs. The RNA component of this complex is considered essential for all organisms except for species of the Aquificaceae family, which contain a protein that does not require the RNA component for tRNA trimming [27], and *Nanoarchaeum equitans*, an obligate symbiont that does not have any detectable RNase P RNA in its reduced genome, nor any detectable RNase P activity [26].

Given the otherwise ubiquitous nature of RNase P RNA, we require detection of this gene as a final quality check of assembled isolate genomes and circularized genomes. However, in the set of circularized CPR genomes in this study, we found that a significant number that lacked RNase P RNA (10/35). Absent a high degree of confidence that these are indeed circular genomes, we would not have noticed this anomaly. We suspected that the RNase P RNA gene was not being detected by the models because the genomes that lacked the gene did not fall into a specific clade and the genes that were detected still had many conserved features of RNase P RNA. To find the missing genes, we reduced the bitscore threshold below the model noise cutoffs when running cmsearch from the Infernal software package [54]. The noise cutoff is the score generally considered to be the score of the highest scoring false positive for that model (Infernal User’s Guide, https://infernal.janelia.org). After reducing the thresholds, we were able to detect the missing RNase P RNAs.

Most of the RNase P RNA genes that we found, even the ones we found initially, required major manual refolding because of the diverged structures with either large extensions of helices (S1 Fig) or missing helices. Many are missing P13, P14, P16, and P17, which is not unusual. However, the RNase P RNA from SRX3307784_bin_224 appears to be missing P12 (Fig 4B), which is highly unusual because this helix is one of the most conserved across the tree of life [40], and it is only known to be missing in *Mycoplasma fermentans* [55] and members of the archaeal family Thermoproteaceae [56]. The closely related genome in this study, ERX2165959_bin_53, is also missing P12 (S2 Fig). Another unusual feature is that approximately two-thirds of the RNase P RNA (23/35) are missing the UGG motif that binds to the CCA in pre-tRNAs. This motif tends to be missing from cyanobacteria and chloroplasts, which may not have the CCA in their pre-tRNAs [57]. Given that cyanobacteria are one of the closer lineages to the CPR in the bacterial tree, the loss of the UGG motif may be related to lineage. A final example of a diverged feature is that the RNase P RNA from SRX1775579_bin_0 appears to be missing nearly the entire P15 helix (S3 Fig). This helix is responsible for establishing binding to pre-tRNAs in bacteria and typically contains the UGG motif, although it is missing from all known RNase P RNA in eukaryotes and some archaea.

Finding these diverged forms of RNase P RNA would not have been possible without having confidence that we had a complete genome. Finding these diverged structures also illustrates that we may find diversity of genes in metagenomics data when we are no longer restricted by what we can culture in the laboratory.

**Fig 4.**
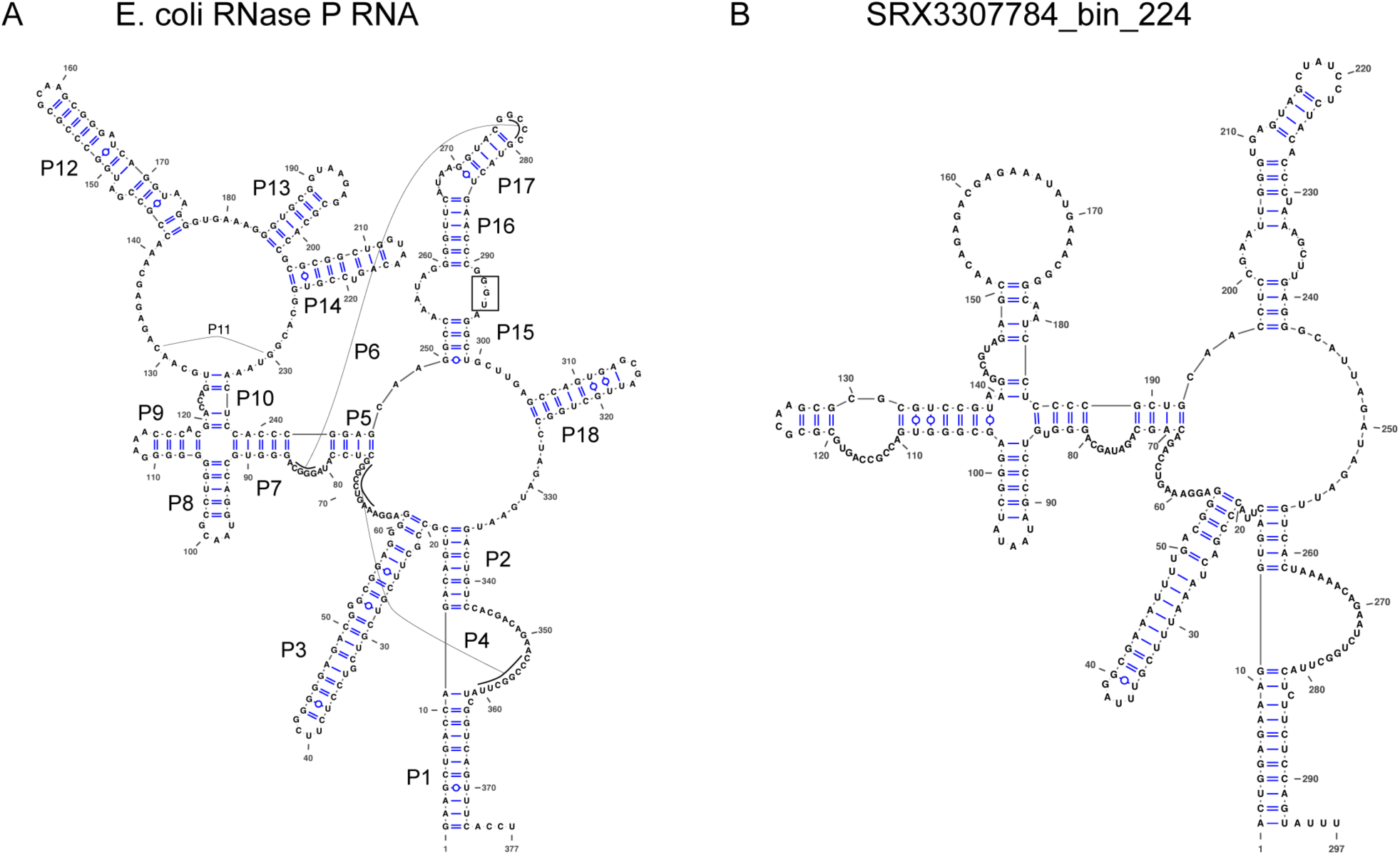
RNase P RNA can have diverged forms in CPR genomes. Structures were drawn using VARNA (Visualization Applet for RNA, http://varna.lri.fr/) [58]. (A) Structure of RNase P RNA from *Escherichia coli K-12* substr. MG1655. Helices P1-P18 are labeled. The “UGG” sequence in the P15 loop that binds to the 3’ end of the pre-tRNAs is highlighted by a box. (B) Putative Structure of RNase P RNA from SRX3307784_bin_224 genome. Note that the P12, P13, P14, and P18 helices are missing, as well as the UGG motif. Although it is not uncommon for P13, P14, and P18 to be missing in various bacteria, it is unusual that P12 is missing. To compare the two structures, large regions of the RNA had to be refolded manually from the original cmsearch prediction.

### Detection and Assembly of Megaphage Genomes

In the process of circularizing genomes, we circularized what we first thought were two novel isolates with small genomes (∼0.5 Mb). However, since one of our standard checks is to run all circular sequences against a full set of Rfam models, these immediately stood out because the only RNAs detected were tRNAs and a tmRNA. Also, GTDB-tk was unable to assign a taxonomy. Cursory BLASTX searches of large regions of the genome yielded only distant hits. Based on this, we decided that they were likely megaplasmids, but have now concluded that they are megaphage based on a recent publication [59].

SRX3024509_bin_4 is an example of one of these putative megaphages. It is 536,059 nt long and codes for 74 tRNA sequences along with one tmRNA. We have seen more than 10 similar - in terms of size and RNA content - megaphages in a variety of environments. They appear to be quite common and if we expanded our size limits, we believe we would find many more. Our method should in theory perform even better on plasmids and viruses than on normal genomes since the former are less likely to have repeats. Extraction of plasmids and viruses from the metagenomes is a matter for future work.

## Discussion

We believe it is crucial to have a substantial collection - on the order of hundreds per phylum - of genomes that approach traditional finished genome standards as closely as possible, such as having a single circular, contiguous sequence with an error rate less than 1 per 100,000 bp [60]. Given that we have not yet succeeded in isolating many of the species found in metagenomics datasets, our focus is on extracting their genomes from environmental metagenomes and enrichments. By checking that assembled genomes are circular and possess all standard known components of genomes - such as RNAs without which life as we know it cannot exist - are present, we gain high confidence that we had nothing but the genome and that the genome was not falsely circularized. We believe that circularity is a top criterion for a high quality assembly, along with checks for misassemblies. We see a clear need for an ongoing curation of collections of genomes. As more circularized genomes are generated from metagenomics data, comparisons will help expose misassemblies and false circularization. For metagenomics data, checking for misassemblies is crucial because they can produce chimeric genomes and lead to erroneous conclusions and information in public genomic databases [46].

During the development of our circularization method, we learned some lessons about when it is the most successful and instances where it will likely fail:

1. Our method works well for small genomes without repeats of significant length. Exact repeats longer than the fragment length remain an issue. If the fragment length is less than the length of a repeat, then it cannot resolve the repeat in the assembly. Once repeats get above the fragment length, the process will - and should - fail.
2. We noted that genomes with rRNA copy numbers greater than one will almost always fail to circularize. We are aware of a single case where MIRA assembled two copies of an SSU on a single contig, so it is possible to do, but it is rare. Binners almost always fail to correctly bin multiple copies of rRNA operons as they end up on shorter contigs with coverage that is a multiple of the single copy stretches. Because we do “digital primer walking”, it is possible to extend a contig to cover a portion of an unbinned contig containing the ribosomal RNA genes While our method will not result in automatic circularization in this case, it can set the stage for further manual curation and possible eventual circularization.
3. Circularization of genomes from metagenomes depends heavily on coverage. All of the genomes we circularized had coverage greater than 29X (Table 2), but it may be possible to circularize a genome with lower coverage. However, in these cases, circularization will generally require manual intervention and we do not know how it would be automated.
4. Circularizing genomes with high GC content is more difficult. This is not particularly surprising given that all of this data was Illumina sequenced and there are known biases against high GC content [61]. It is also possible that genomes with high GC content are larger and hence more difficult to succeed with circularization.

Even for bins that did not achieve circularity, if the bin didn’t become Chaos, we suggest that it likely still improved and, in some cases, significantly in terms of the length of contigs and number of misassemblies. MIRA is conservative in extending contigs, so in some cases we believe that with some manual intervention these bins can be circularized.

We are well aware that circularity is not sufficient in and of itself to account for all of the genetic material of a microbe, *i.e*., multiple chromosomes, plasmids, etc. may comprise a genome and not just one chromosome. As long read technology such as PacBio and Nanopore become more feasible for metagenomics, assembling microbial genomes will also become easier [18,23]. In addition, long read technologies are starting to yield methylation patterns that can be used to associate multiple replicons with each other, so it may be possible to resolve if there are multiple chromosomes and plasmids in a genome [62].

We have at least another 100 CPR genomes that are close to being circularized and we believe will only require only minor curation to achieve full circularization. We will release these genomes in future publications. By generating a set of high quality reference genomes, we intend to use these as scaffolds for future assemblies and comparative genomics. Since these genomes are complete, we can also begin to study the genomic context of genes, such as synteny and operons. As we demonstrated with our investigation into RNase P RNA, the study of non-coding RNAs in the intergenic regions of these genomes is also now more feasible than ever. Finally, we also plan to link the 16S to the functional potential that we can see in the genome, allowing us to glean more information from 16S studies in terms of metabolic inference. As we continue to generate high quality complete genomes from metagenomics data, we will be able to more accurately analyze the functional potential of microorganisms that we cannot yet culture.

## Materials and Methods

### Metagenomics Datasets

We used datasets from the NCBI Sequence Read Archive (https://www.ncbi.nlm.nih.gov/sra). Accession numbers are listed in Table 1. In addition, we obtained the SURF datasets from Lily Momper [35].

### Read Processing Assembly

Metagenomic reads were preprocessed using BBtools version 38.60 to remove Illumina adapters, perform quality filtering and trimming, and remove PhiX174 spike-ins. We are not aware of any published papers documenting these tools. However, it is a standard tool suite developed at the Department of Energy Joint Genome Institute (JGI) and it is documented at https://jgi.doe.gov/data-and-tools/bbtools/. Processing was done in two passes. First bbduk.sh ran with parameters *ktrim=r k=23 mink=11 hdist=1 ref=adapters.fa tbo tpe 2*. This was to remove any remaining Illumina adapters given in adapters.fa (standard Illumina adapters). Then bbduk.sh was run again with parameters *bf1 k=27 hdist=1 qtrim=rl trimq=17 cardinality=t ref=phix174_Illumina.fa*. This was to perform quality filtering and trimming as well as remove Illumina PhiX174 spike ins given in the file phix174_Illumina.fa.

### Genome Assembly and Classification

Assembly was performed using SPAdes version 3.13.0 [36] with parameters *--meta -k 21,33,55,77,99,127*. Following assembly, from BWA version 0.7.17-r1188 [63], we used the BWA-MEM algorithm with default parameters to map the reads to the set of contigs produced by the assembly. We did this to obtain the BAM file required by MetaBAT 2 version 2.0 [37]. We used MetaBAT 2 with parameters *--unbinned --minContig 1500 --maxEdges 500* to bin the contigs. The iterative assemblies were performed using MIRA 5.0rc1 [7]. The parameters set were *-NW:cac=warn, -CO:fnic=yes -AS:nop=6:sdlpo=no -KS:fenn=0.3*. We used Pilon version 1.23 [42] with default parameters to run final read coherence checks and clean up issues created by the circularization. Taxonomic classification was generated using GTDB-Tk version 0.3.3 [38].

We carried out all of the work using standard Haswell architectures with 20 cores and 256 GB of main memory. SPAdes is generally memory limited and that is where the high point of memory use occurred. Most of the iterative binning work is possible on a standard desktop or even a laptop with 32 GB of memory as long as the coverage of the candidate genomes doesn’t exceed ∼100X.

### Gene Annotation

All of the RNA annotations were generated by Infernal 1.1.2 [54] using *cmsearch* with parameters *--notextw --cut_tc*. We also used in-house scripts to handle RNA clan processing [64]. We used RFAM version 14.1 for the models except when we used SSU-ALIGN (see Phylogenetic Tree section below) which uses built-in custom models. For RNase P RNA, we reduced the required bit score threshold 5 using the bacterial Class A model (RF00010) to find the diverged forms. Gene calling was done using Prodigal version 2.6.3 [65]. We used *prodigal* with parameters *-n -p single*.

### Phylogenetic Tree

A tree was constructed from a structural alignment of the 16S genes generated by SSU-ALIGN version 0.1.1 with default parameters [66,67]. Some 16S genes required manual folding and adjustments to correct the structural alignment. We used IQ-TREE version 2.0-rc1 [45] to generate the tree via the web server at Los Alamos National Laboratory.

### Circularization code

The code and a tutorial of this method is available on Github (https://github.com/lmlui/Jorg).

## Data Availability

Assembled genomes from this study were deposited at NCBI under BioProject accession numbers PRJNA416593, PRJEB22302, PRJNA640834, PRJNA640835, PRJNA640837, PRJNA640839, PRJNA640841, PRJNA640844-PRJNA640846, PRJNA640848, PRJNA640849, PRJNA640851-PRJNA640855, PRJNA640857, PRJNA640860-PRJNA640863, PRJNA640871-PRJNA640876, PRJNA640878, PRJNA640925, PRJNA640926, PRJNA640929, PRJNA641932, PRJNA641935, and PRJNA641936, and PRJNA641940.

## Acknowledgements

This work was funded by ENIGMA, a Science Focus Area Program supported by the U. S. Department of Energy, Office of Science, Office of Biological and Environmental Research, Genomics Sciences Program: Foundational Science; this research also used resources of the Joint Genome Institute (JGI) and the National Energy Research Scientific Computing Center (NERSC), U.S. Department of Energy Office of Science User Facilities; all managed by Lawrence Berkeley National Laboratory under Contract No. DE-AC02-05CH11231. The authors would also like to thank members of the ENIGMA Science Focus Area as well as Heidi Smith, Patricia Chan, and Norman Pace for helpful feedback and suggestions. We would also like to thank Sean Jungbluth and Lily Momper for giving us access to their datasets.

## Supplementary figures

**S1 Fig.**
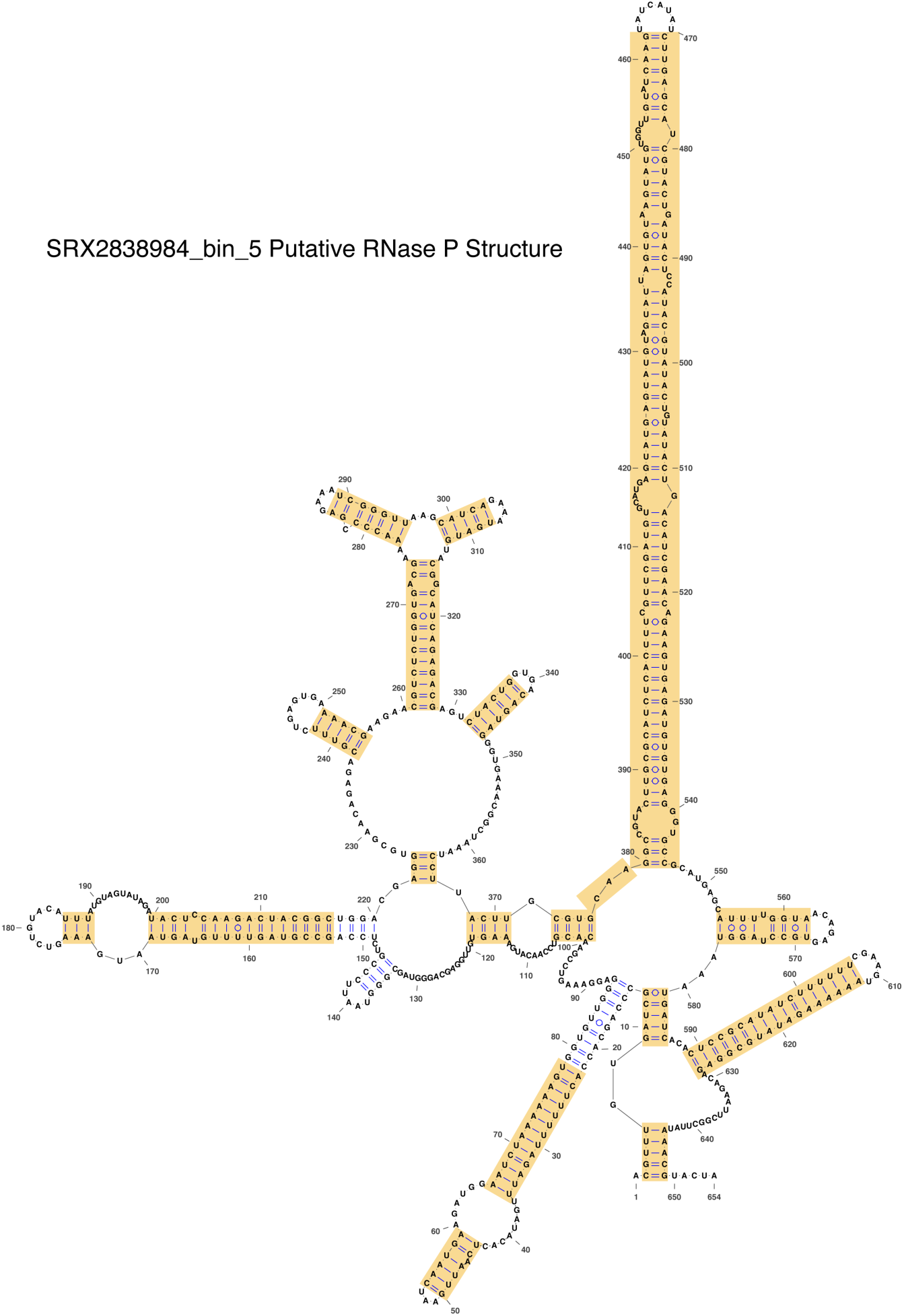
Putative RNase P RNA structure of SRX2838984_bin_5. This RNase P RNA appears to have an extended P15 helix compared to typical RNase P RNA (see the *E. coli* RNase P RNA structure in Fig 4A of the main text). Yellow highlights indicate the portions of the RNA that had to be refolded manually. This amount of refolding was not unusual for the RNase P RNAs found in this study.

**S2 Fig.**
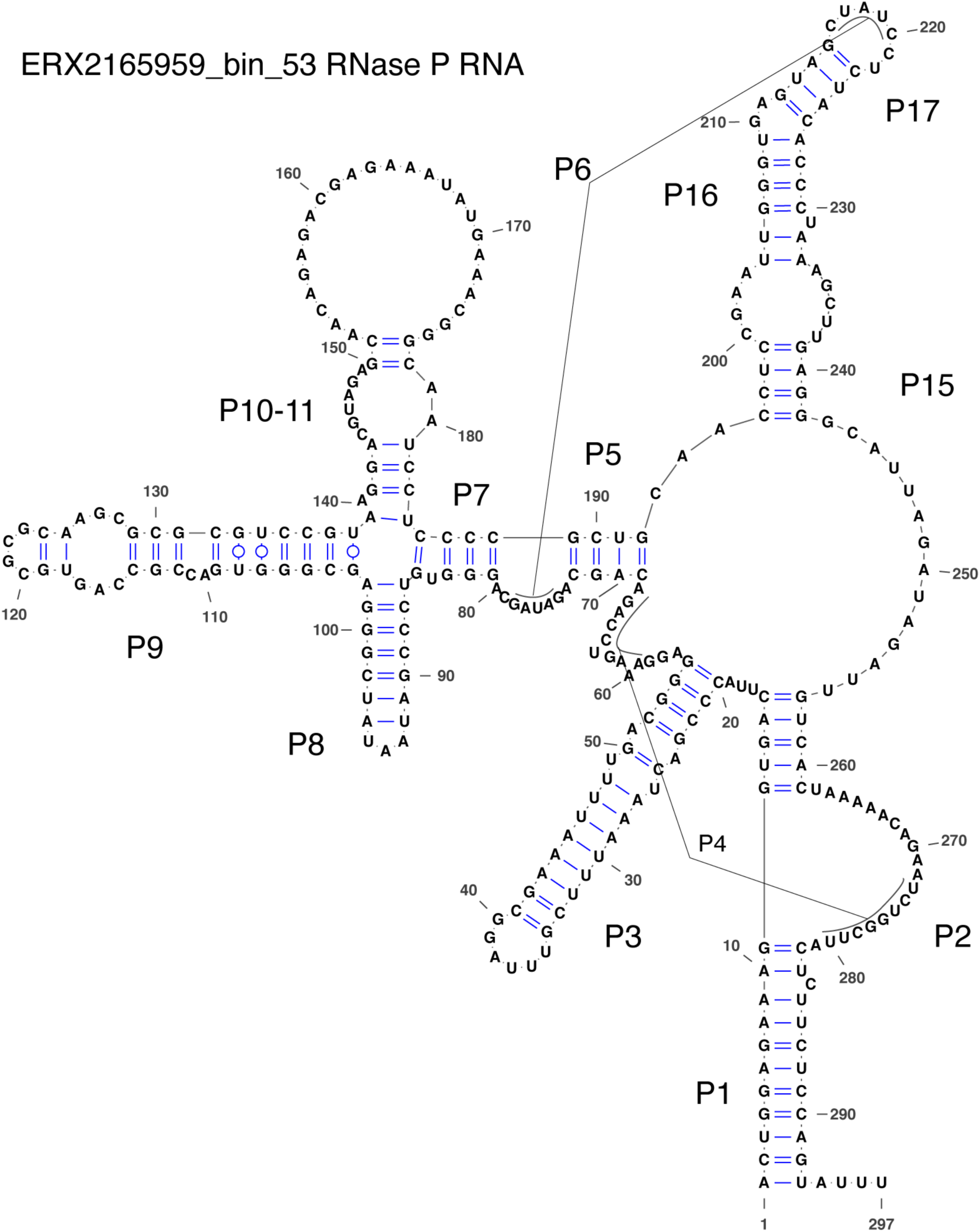
Putative RNase P RNA structure of ERX2165959_bin_53. This structure is missing P12, P13, P14, and P18. It is not unusual to be missing these helices, except for P12 which is found in nearly all RNase P RNA structures.

**S3 Fig.**
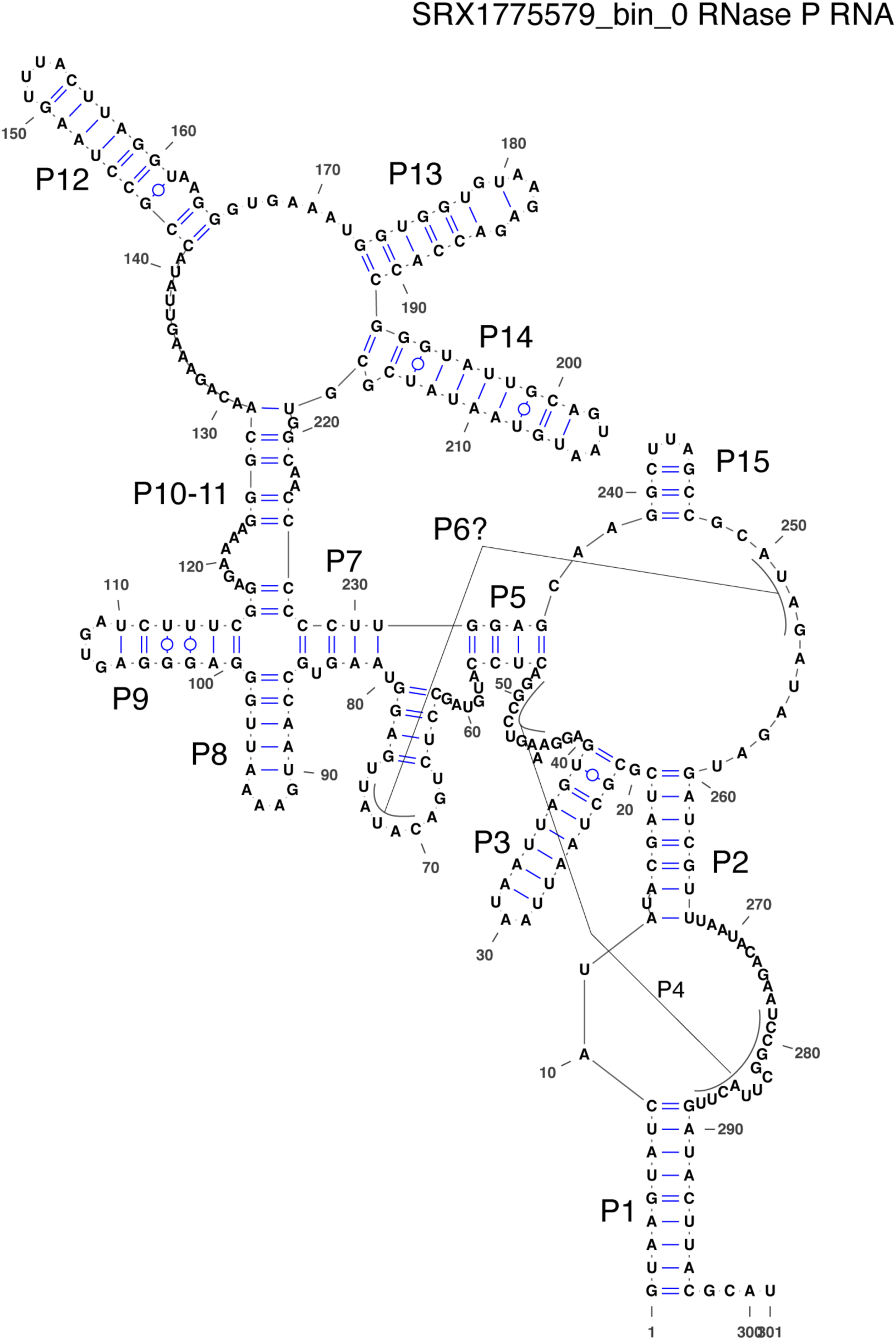
Putative RNase P RNA structure of SRX1775579_bin_0. This structure appears to be missing most of the P15 helix.

